## Supplementary Information for "Biomolecular condensates modulate membrane lipid packing and hydration"

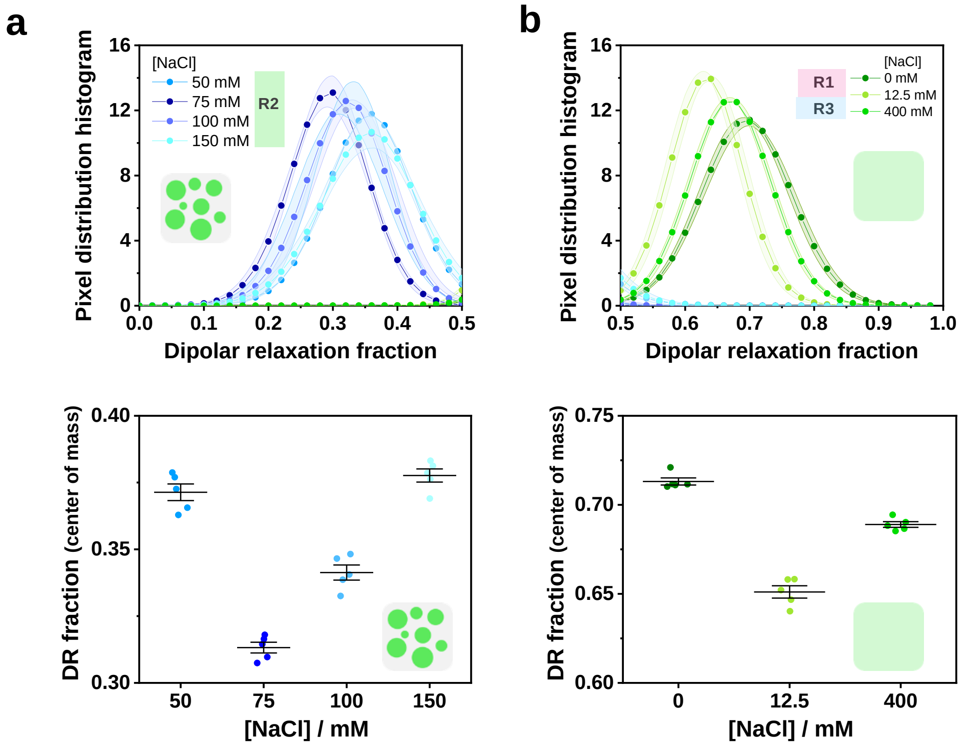


Supplementary Figure 1: **ACDAN senses a different nano-environment in the homogeneous (R1 and R3) and condensate phases (R2).** Zoomed plots of those shown in Fig 2d,e. The upper panels show the histograms, and the bottom panels show the center of mass for **a.** Glycinin in the condensate phase (R2), and **b.** Glycinin in the homogeneous phase (R1 and R3), at the indicated NaCl concentrations; for regions R1, R2 and R3, see the phase diagram in Fig. 2a. Histograms and center of mass are shown as mean±SD, n=5 per condition. Data for panels a-b are provided as a Source Data file.

**Supplementary Table 1** Full width at half maximum (FWHM) input values in $cm^{-1}$ and their physically plausible ranges expected for each type of secondary structure as measured with ATR-FTIR.[1, 2]

| Secondary structure component | | FWHM input ($cm^{-1}$) | Lower limit ($cm^{-1}$) | Upper limit ($cm^{-1}$) |
|---|---|---|---|---|
| *β-sheet* | *High wavenumber component (1670 – 1680)* | 9 | 8 | 11 |
| | *Low wavenumber component (1610 – 1640)* | 22 | 11 | 33 |
| *Random (1640 – 1650)* | | 55 | 50 | 60 |
| *α-helix (1650 – 1660)* | | 20 | 5 | 30 |
| *Turns (1660 – 1670; 1680 – 1700)* | | 20 | 5 | 30 |

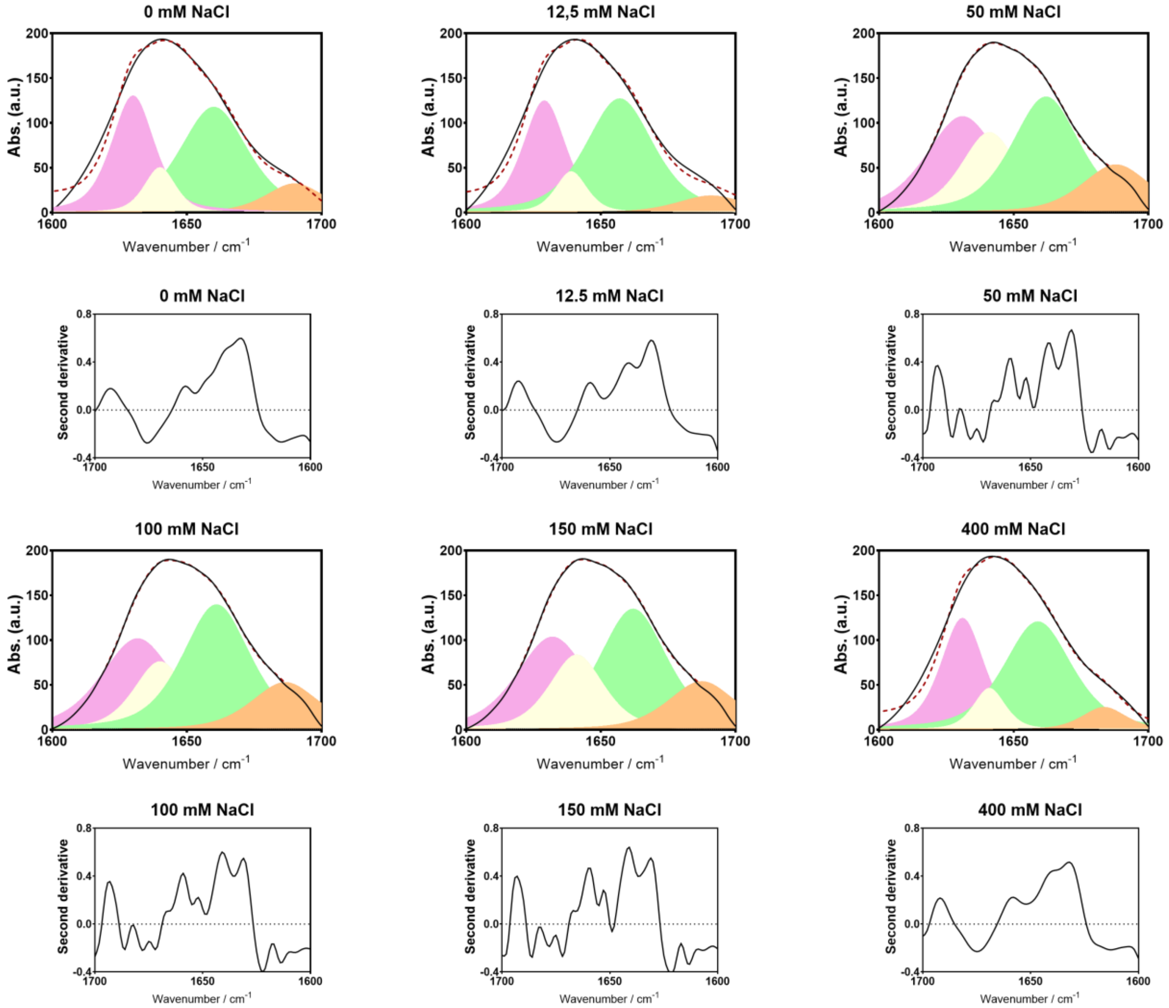


Supplementary Figure 2: **ATR-FTIR analysis.** ATR-FTIR Amide I band of glycinin (10 mg/ml) for the different salinity conditions (solid lines) and best fits for the band components (red dashed lines). The components correspond to the colors in Table S1: pink for β-sheet, yellow for random-coil, orange for turns, and green for α-helix. The respective second derivative of the Amide I spectrum is shown below each case. Data for all panels are provided as a Source Data file.

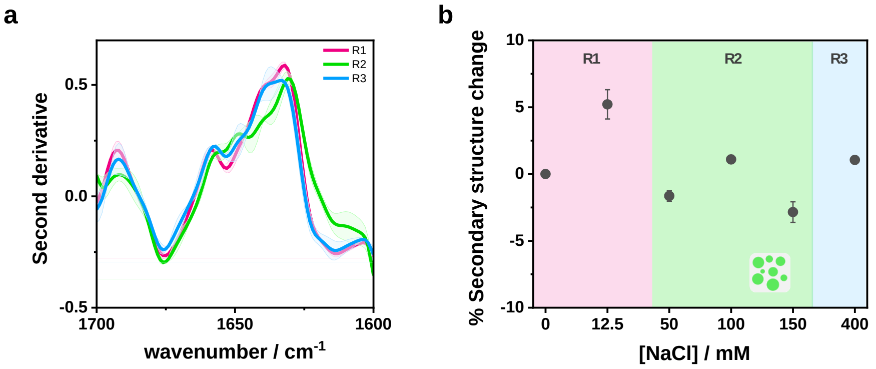


Supplementary Figure 3: **Glycinin secondary structure content. a.** Examples of the second derivative of the Amide I band of glycinin at the different regions of the phase diagram (see Fig. 2a): R1 (0 mM NaCl), R2 (100 mM NaCl), R3 (400 mM NaCl). **b.** Change in the β-sheet content compared to the initial values for the protein in water. The changes are less pronounced, and there is no clear tendency as in the case of α-helix and random+turns as depicted in Figure 3. Data for panels a-b are provided as a Source Data file.

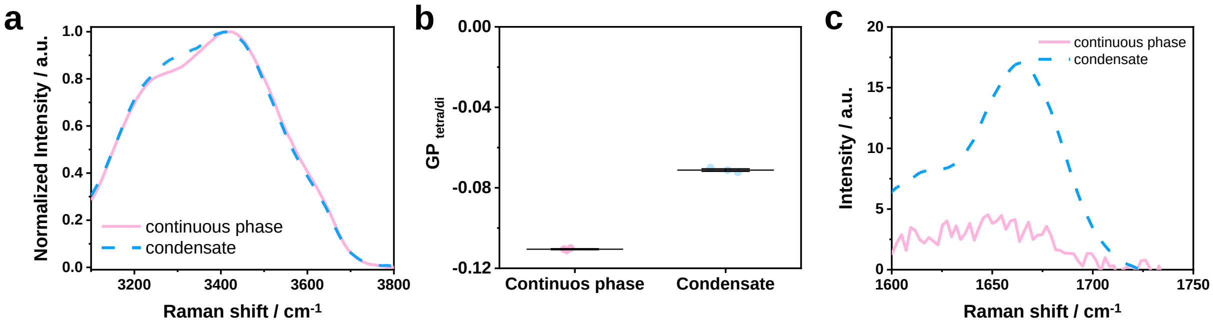


Supplementary Figure 4: **Raman spectra analysis. a.** Examples of the water band for the continuous and condensate phase at 100 mM NaCl. The signal for the condensate phase appears shifted towards lower wavenumbers, and the band intensity at and around 3225 cm$^{-1}$ is increased compared to the continuous phase. **b.** GP function (eq. 9) quantifies the spectral shift and intensity change in a. A higher GP value indicates an increase in tetra-coordinated water. Individual points are shown as circles and the lines indicate mean±SD. **c.** Raman spectra of the Amide I region at 100 mM NaCl, show that the protein signal is high in the condensates and negligible in the protein-depleted phase. Data for panels a-c are provided as a Source Data file.

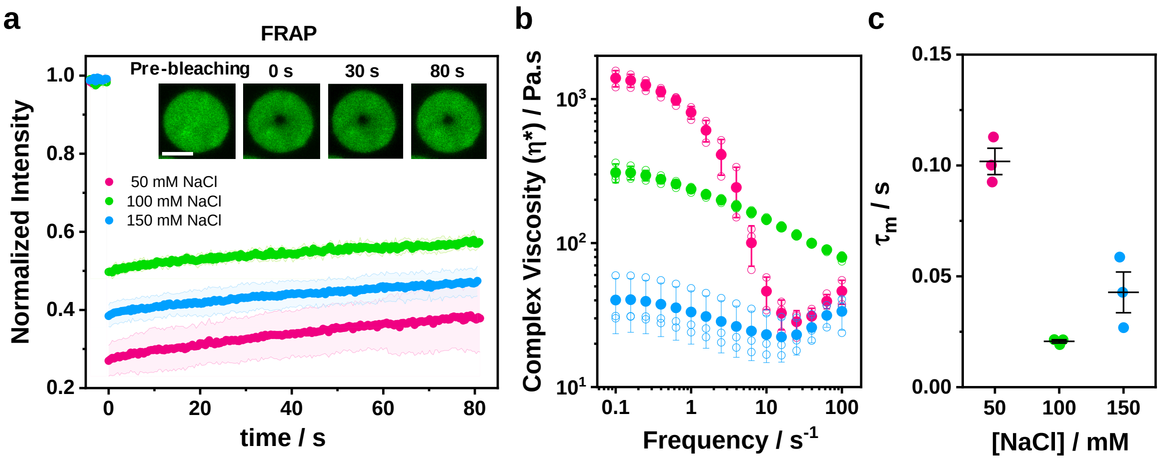


Supplementary Figure 5: **Condensates material properties. a.** Fluorescence recovery after photo bleaching (FRAP) for glycinin condensates show negligible recovery, reflecting the high viscosity of these condensates, as previously reported[3]. Glycinin concentration is 10 mg/mL. Data show mean values and the shadowed area corresponds to the standard deviation (n=4 per condition). The insets show an example of a condensate at 100 mM NaCl before and after bleaching at the indicated times. Scale bar is 5 µm. **b.** Complex viscosity ($\eta^*$) vs frequency obtained by rheology measurements for the protein-rich (condensate) phase at different NaCl concentrations: 50 mM (pink), 100 mM (green), 150 mM (blue). Individual data points are shown as open circles and filled circles are mean±SD (n=3 per condition). **c.** Terminal relaxation time ($\tau_m$) for the condensates at different NaCl concentrations. Individual data points are shown as circles, lines represent the mean±SD. Data for panels a-c are provided as a Source Data file.

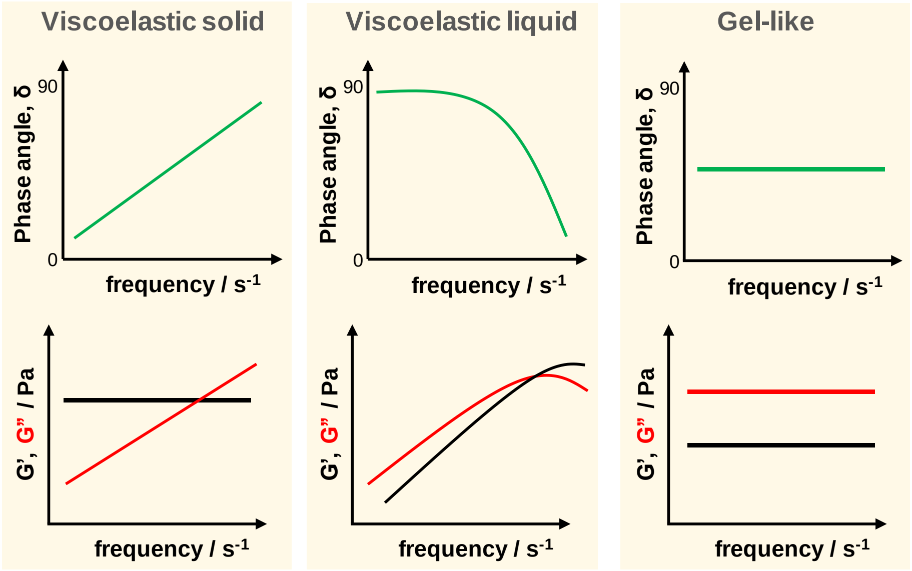


Supplementary Figure 6: **Rheology measurements interpretation.** The sketches exemplify the typical responses of phase angle ($\delta$) and storage ($G'$) and loss ($G''$) moduli vs frequency for different types of materials. Adapted from reference [4].

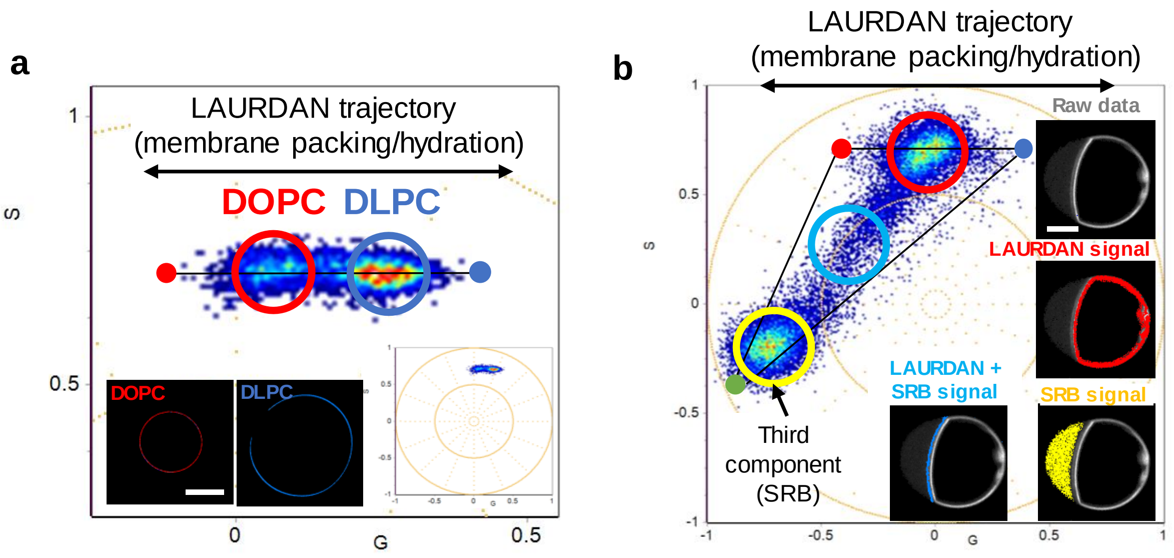


Supplementary Figure 7: **LAURDAN fluorescence in different membranes describes a linear trajectory between various hydration states. a-b.** Proof-of-principle experiments showing the response of LAURDAN in membranes and how it changes when a third component is incorporated. **a.** DOPC and DLPC membranes labeled with LAURDAN (0.5 mol%) display a linear trajectory in the phasor plot, corresponding to different hydration and packing states. **b.** Fluorescence signal from probes other than LAURDAN do not appear along the linear trajectory (as displayed in panel a). This is exemplified with fluorescence signal from the condensates when labeled with the water-soluble dye Sulforhodamine B (SRB). In the phasor plot, it appears as a third component allowing its clear identification and separation of the signals, as exemplified here. For cases like this, a three cursor analysis would be required[5] to unmix the signals and quantify the measured processes, in a similar way as the two-cursor analysis employed in this work.

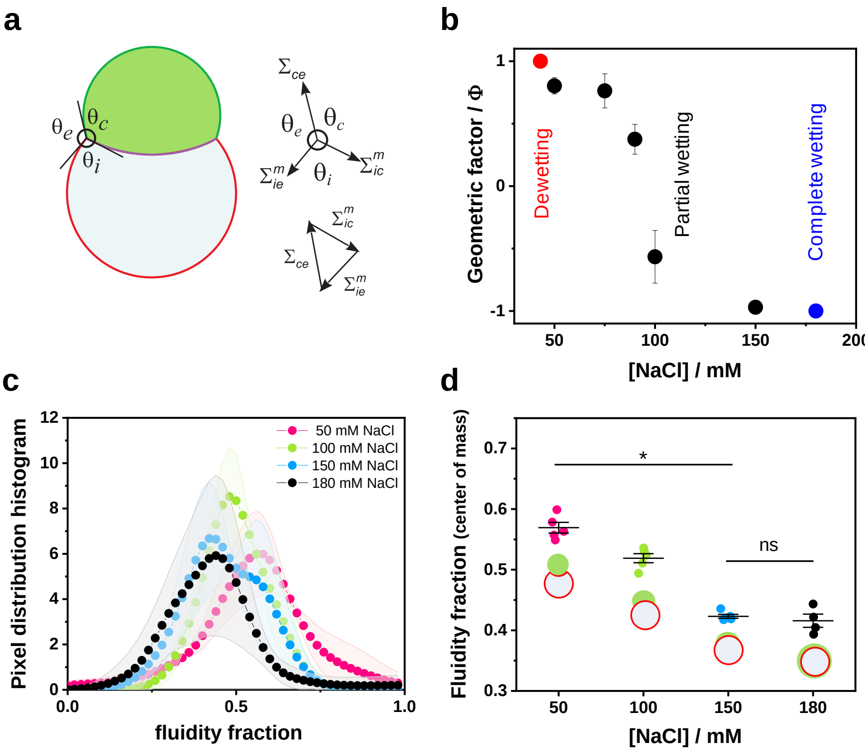


Supplementary Figure 8: **Geometric factor and interaction affinity and their relation to fluidity. a.** A sketch showing the three contact angles between the two membrane segments and droplet interface with the external solution. The contact angles and the respective tensions are related via the force balance triangle shown in the right. By measuring the angles with optical microscopy, the geometric factor ($\Phi$) can be obtained as: $\Phi = (\sin\theta_e - \sin\theta_c)/\sin\theta_i$, for details see reference 6. This dimensionless factor is independent of the relative sizes of the droplet and vesicle and is determined by the material properties of the condensate and the membrane. The geometric factor provides an indirect measurement of the affinity contrast between the condensate and the membrane with respect to the external solution. **b.** Geometric factor for glycinin condensates in contact with vesicles at different NaCl concentrations. The system undergoes two wetting transitions, from dewetting ([NaCl]=43mM) to partial wetting (43>[NaCl]>180) to complete wetting ([NaCl]=180 mM). **c.** Fluidity fraction histograms for vesicles in contact with glycinin condensates at the indicated NaCl concentrations. Data show mean±SD (n=5). **d.** Center of mass distribution for the histograms shown in c. There are not significant differences between the salt conditions of 150 and 180 mM NaCl. Individual data points are shown, the lines indicate mean±SD. Note that panels c and d show the same data as in Figure 6c, except for the composition of 180 mM NaCl. Panels a and b are adapted from reference 6. Data for panels c-d are provided as a Source Data file.

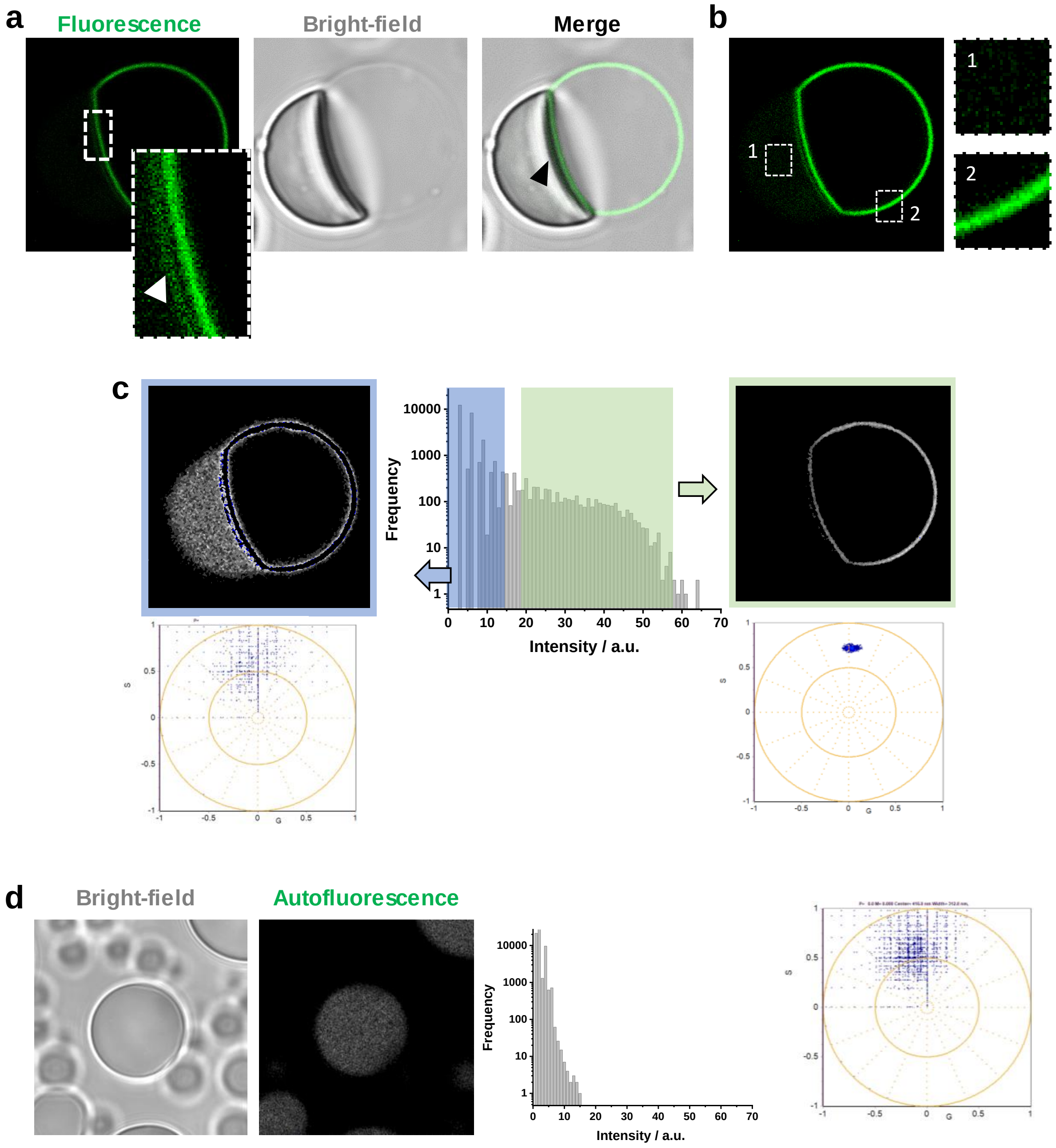


Supplementary Figure 9: **Protein autofluorescence is negligible and does not interfere with LAURDAN measurements. a.** The image displayed in Figure 6b of a DOPC GUV labeled with LAURDAN in contact with an unlabeled glycinin condensate at 100 mM NaCl, shows a double fluorescent line at the interface between the membrane and the condensate (see zoomed panel). This is due to an optical effect arising from the high refractive index of the condensates. **b.** LAURDAN fluorescence comes exclusively from the membrane, but when contrast is increased weak signal from protein autofluorescence can be detected from the condensate as exemplified in the zoomed panels. **c.** When analyzing the pixel intensity distribution with the phasor approach we can see that the autofluorescence contribution corresponds to low intensity pixels (shaded in blue), and appear as noise in the spectral phasor plot. We can eliminate these pixels and only analyze the pixels coming from the membrane (shaded in green), that appear as a coherent cloud in the phasor plot shown below the image. Note that we also eliminate high intensity and saturated pixels. **d.** The cutoff intensity in panel c is selected from measurements on condensates autofluorescence in the absence of GUVs and with the same setup used throughout the work (see Methods). Similar to panel c, the autofluorescence signal is very low and appears as noise in the phasor plot. Data for panels c-d are provided as a Source Data file.

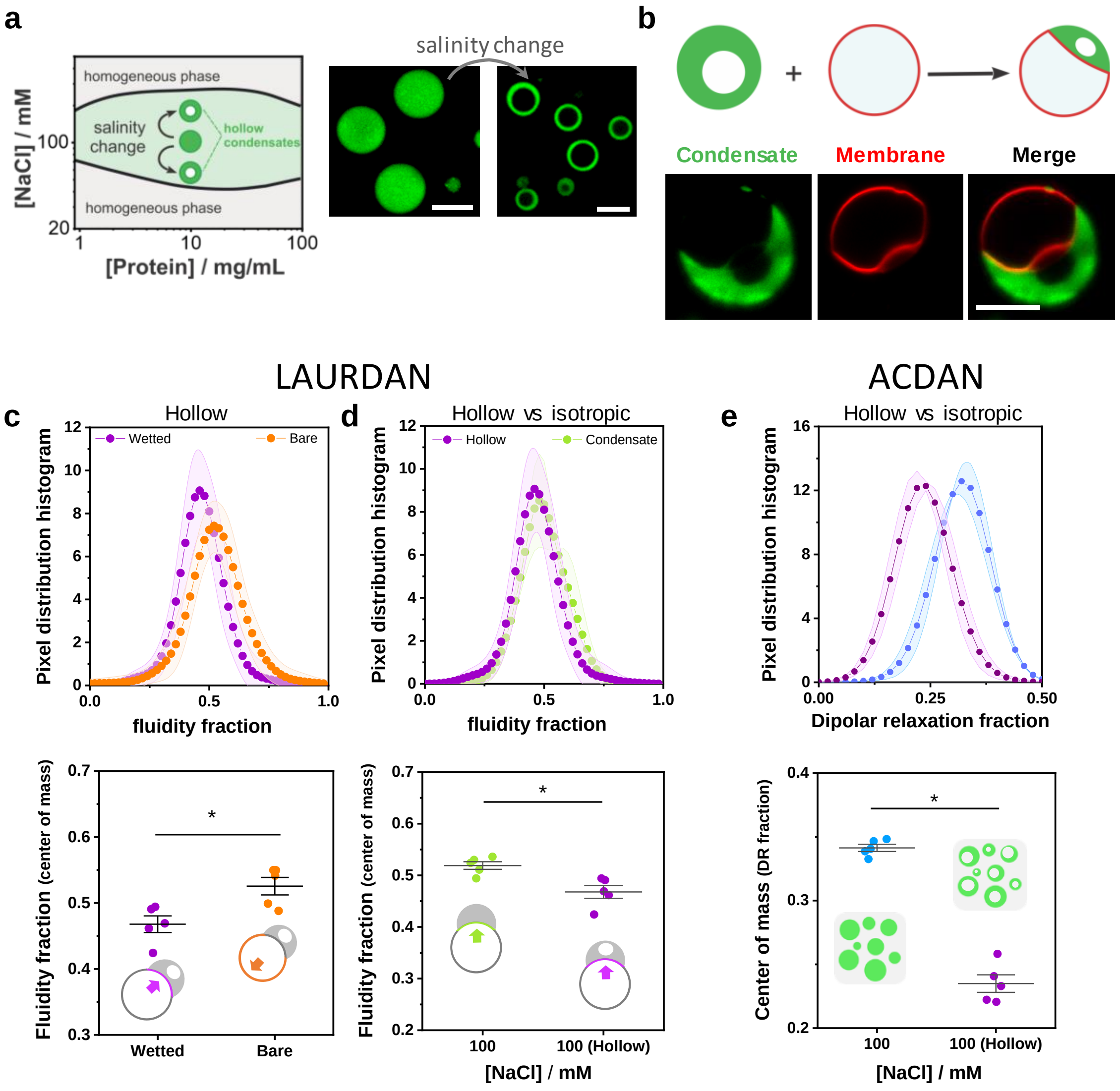


Supplementary Figure 10: **Hollow condensates effect on membrane packing. a.** Hollow condensates can be formed exposing (isotropic) condensates to a sudden change in salinity to trigger phase separation within them, as indicated in the phase diagram sketch. This leads to the formation of a protein-poor phase ("hollow" void) surrounded by the protein-rich phase[3, 6]. The images show glycinin condensates at 100 mM NaCl that become hollow after diluting the sample to 50 mM NaCl. **b.** Hollow condensates in contact with GUVs can wet and mold the membrane in a similar way as isotropic condensates. The confocal images are an example of a hollow condensate (green) wetting a GUV (red) at 150 mM NaCl. **c-e.** Histograms (upper panels) and center of mass (lower panels) obtained through hyperspectral imaging and phasor analysis of LAURDAN and ACDAN at 100 mM NaCl. Individual points are shown as circles and lines are mean±SD, n=5. **c.** Comparison between the bare and wetted segments of vesicles in contact with hollow condensates at 100 mM NaCl (final concentration in both cases). Similar to the behavior of isotropic condensates, the membrane wetted by the hollow condensate presents an increased packing compared to the bare membrane. Note that the fluidity fraction values obtained for the bare membrane correspond to those obtained for the bare membrane of GUVs in contact with isotropic condensates (compare to Fig. 6c). **d.** Comparison between the fluidity fraction for membranes in contact with isotropic or hollow condensates at 100 mM NaCl. The hollow condensates generate an increased membrane packing compared to isotropic ones at the same salinity. **e.** ACDAN shows a very different response for hollow condensates compared to isotropic ones at the same salinity. The lower dipolar relaxation observed for hollow condensates could imply differences in the protein secondary structure and hydrogen bonding. Figures a and b are adapted from reference [6]. All scale bars are 10 µm. Data for panels c-e are provided as a Source Data file.

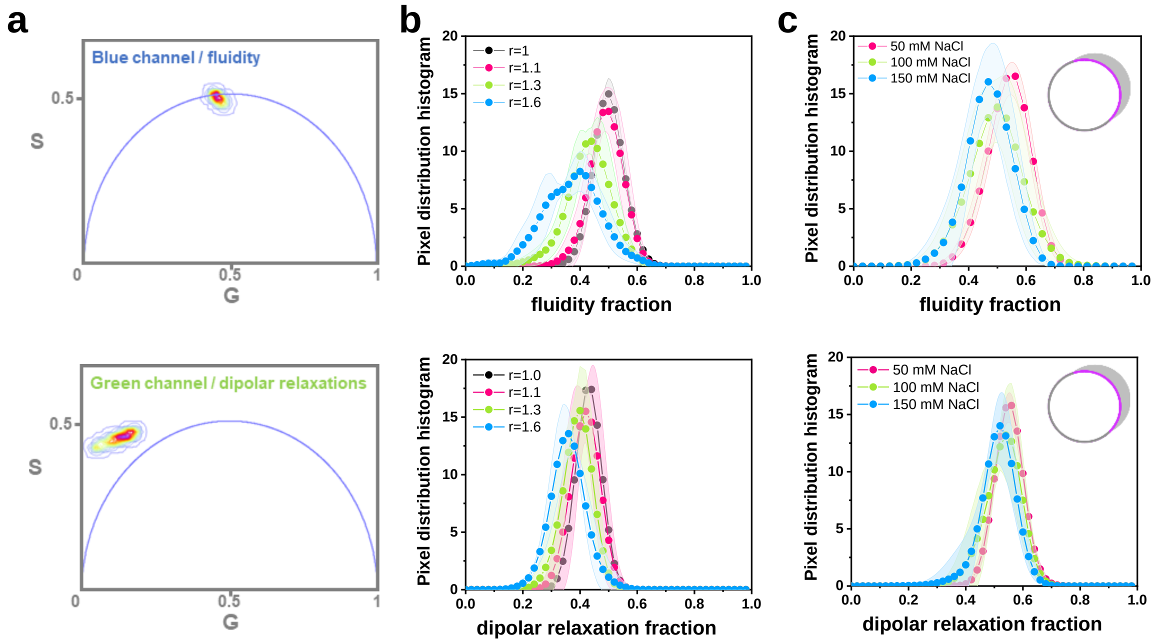


Supplementary Figure 11: **FLIM two-channel phasor analysis.** Using a blue and a green filter, LAURDAN FLIM phasor plots allow discrimination of the fluidity and water dipolar relaxation changes occurring in the system. **a.** Examples of the pixel clouds in the blue (top panel) and green (bottom panel) channels for measuring the polarity/fluidity and dipolar relaxation changes, respectively. Note that, as expected, for the dipolar relaxation channel, the pixel cloud is located outside the universal circle; this is a signature of processes occurring while the probe is in the excited state[7, 8]. Fluidity (top) and dipolar relaxation (bottom) histograms for the ATPS **(b)** and GUVs in contact with glycinin condensates systems **(c).** For both systems, an increase in wetting correlates with increased lipid packing and a decreased dipolar relaxation. The sketches indicate the part of the membrane analyzed for the glycinin systems. Curves are shown as mean values (circles) ± SD (shadowed area), n=5. Data for panels b-c are provided as a Source Data file.

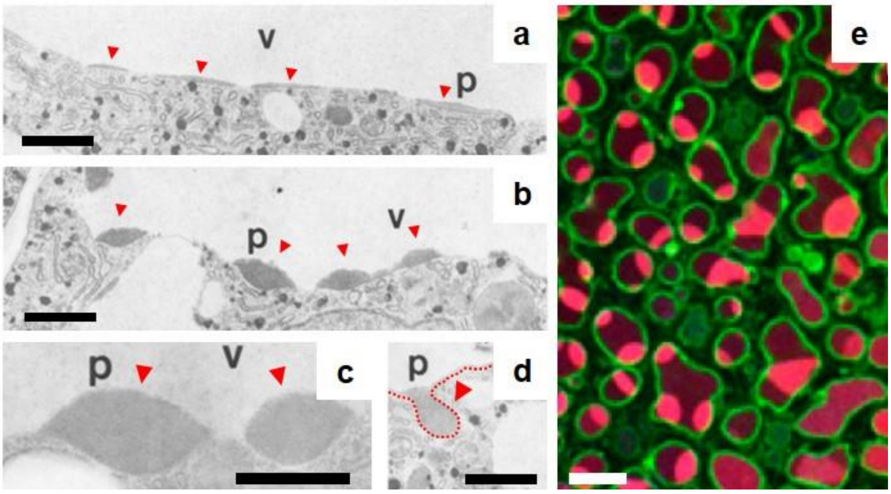


Supplementary Figure 12. **Wetting and remodeling of plant vacuolar membranes by storage protein condensates. a-d.** Electron micrographs of storage parenchyma cells of soybean cotyledons during development (p=protein droplets, v=vacuole): (a) protein droplets (red arrowheads) spread on the vacuolar membrane, (b, c) partially wet it, imposing additional curvature in the contact regions, and (d) forming “protein pockets” as finger-like structures (the red dotted line highlights the membrane contour) protruding towards the cytoplasm as a step prior to protein body formation. **e**. Confocal microscopy image of protein storage vacuoles in Arabidopsis thaliana embryo cells: the protein droplets (red) wet the vacuolar membrane (green). Scale bars in (a-d) are 1 µm and in (e) 10 µm. Images (a-e) were adapted from reference [9] with permission from SNCSC and image (f) from reference 10.

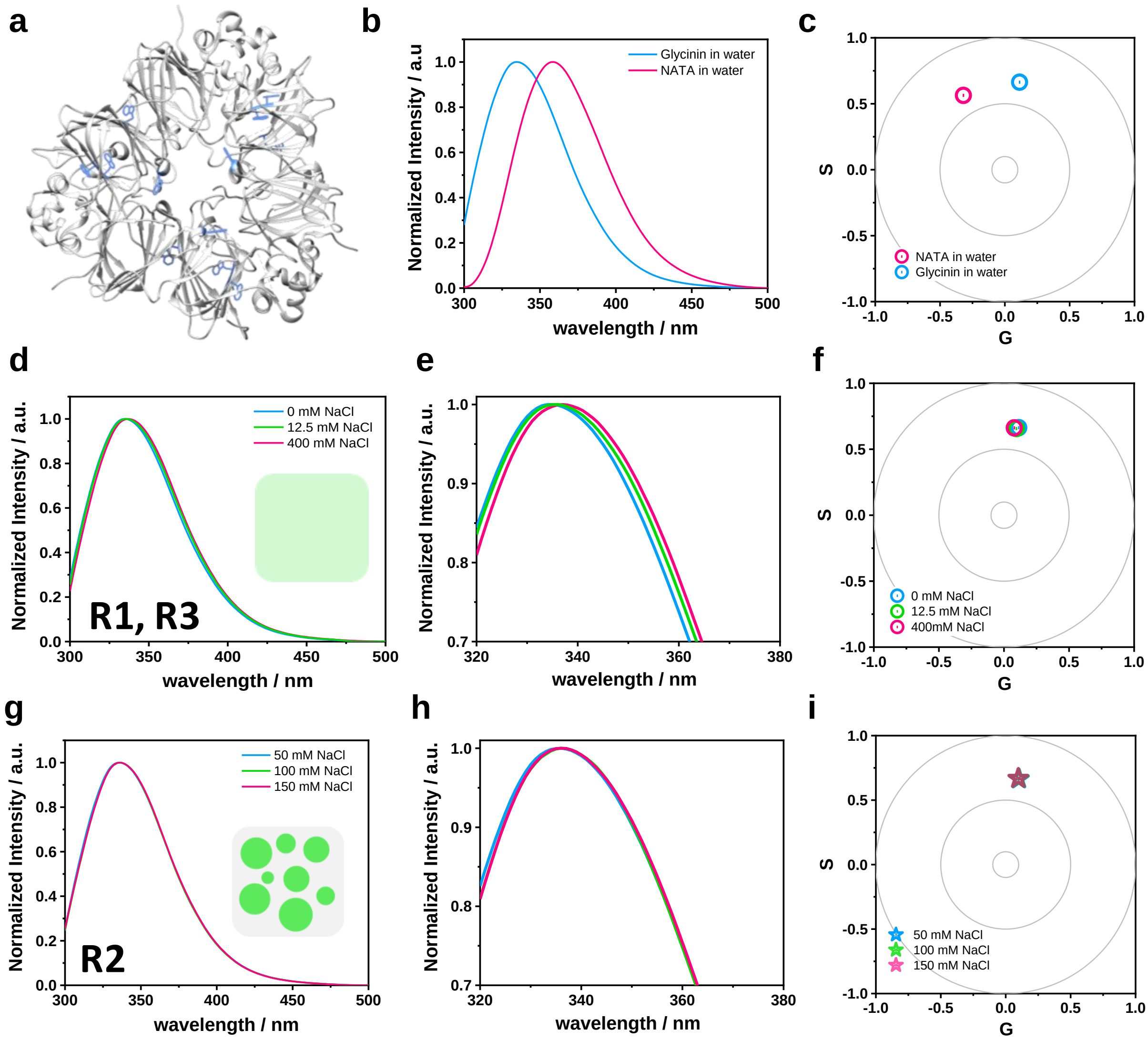


Supplementary Figure 13: **Glycinin autofluorescence is unsuitable for detecting structural/environmental changes. a.** Glycinin crystallographic structure[11] with the tryptophan residues highlighted in blue. For clarity, only one trimer of the glycinin hexamer is shown. Glycinin contains 24 tryptophan residues, most of which are oriented toward the protein's interior. **b.** Fluorescence emission spectra of N-Acetyl-L-tryptophanamide (NATA) in water and glycinin in water. NATA is a standard for tryptophan emission in water, representing the fluorescence of a tryptophan residue completely exposed to the solvent. The excitation wavelength was 280 nm. It can be seen that the spectrum for glycinin appears blue-shifted with a maximum near 335 nm, while NATA maximum is near 360 nm. **c.** Spectral phasor graph of the spectra shown in (b). The spectral shift is considerable and indicates that glycinin's tryptophan residues environment is highly hydrophobic. **d.** Glycinin tryptophan spectra for the homogeneous phase (R1, R3) at the indicated NaCl concentrations. **e.** A zoom of the spectra shown in d reveals a small shift. **f.** Spectral phasor graphs of the spectra shown in (d, e). It can be seen clearly that the shift is minimum compared to the difference shown in c. **g.** Glycinin tryptophan spectra for the condensate phase (R2) at the indicated NaCl concentrations. **h.** Zoom of the spectra shown in (g). **i.** Spectral phasor graphs of the spectra shown in (g, h). Again, the changes are negligible, demonstrating that the sensitivity of protein autofluorescence is not suitable for addressing the environmental and structural changes occurring at different conditions. For this reason, it is necessary to use an external reporter to address such changes. Data for panels b-i are provided as a Source Data file.

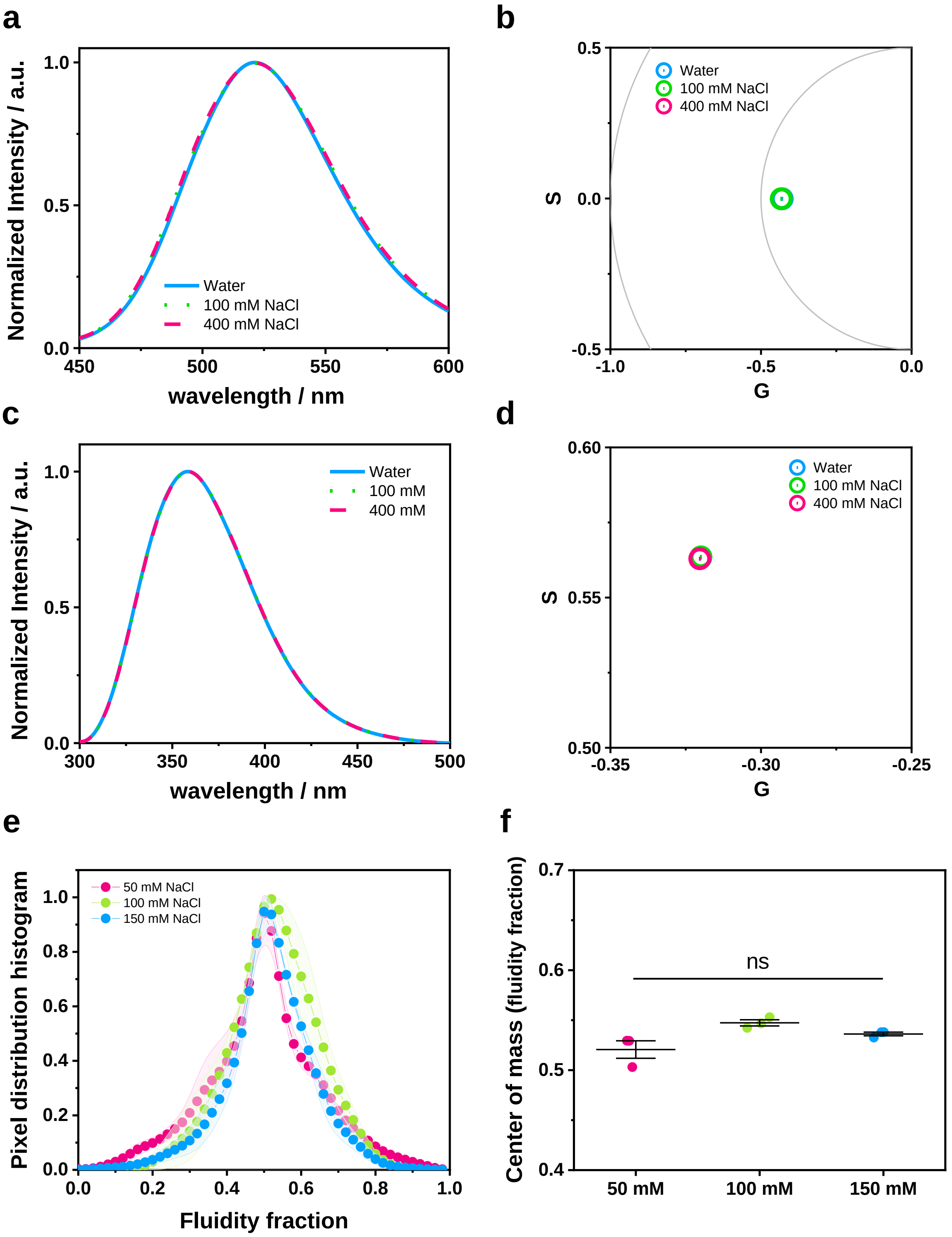


Supplementary Figure 14: **Sodium chloride concentration does not affect dye fluorescence in the studied range. a.** Fluorescence spectra of ACDAN in the indicated conditions (ex. 360 nm). **b.** Spectral phasor analysis of ACDAN spectra shown in (a). **c.** Fluorescence spectra of NATA in the indicated conditions (ex. 280 nm). **d.** Spectral phasor analysis of NATA spectra shown in (c). **e.** DOPC GUVs LAURDAN fluorescence histograms obtained from the spectral phasor plot. **f.** The center of mass of the histograms shown in (e). Individual measurements are shown as circles and lines represent mean±SD. There are no significant differences between the conditions. Data for panels a-f are provided as a Source Data file.
